# Peak alpha frequency is not significantly altered by five days of experimental pain and repetitive transcranial stimulation of the left dorsolateral prefrontal cortex

**DOI:** 10.1101/2024.06.14.599003

**Authors:** Samantha K. Millard, Alan K.I. Chiang, Nahian Chowdhury, Wei-Ju Chang, Andrew J. Furman, Enrico De Martino, Thomas Graven-Nielsen, Siobhan M. Schabrun, David A. Seminowicz

## Abstract

Repetitive transcranial magnetic stimulation (rTMS) holds promise as a non-invasive pain treatment. Given the link between individual peak alpha frequency (PAF) of resting-state electroencephalographic recordings and pain sensitivity, and the potential for rTMS to modulate PAF, we investigated these relationships through a secondary analysis of established rTMS-induced analgesia in an experimental model of sustained muscle pain.

In a randomised, single-blind, sham-controlled experiment, 30 healthy adults underwent either active (n=15) or sham (n=15) high-frequency rTMS (20 min) to the left dorsolateral prefrontal cortex for five consecutive days following induction of sustained experimental pain by nerve growth factor (NGF) injected into the right extensor carpi radialis brevis muscle. The pain intensity was assessed daily for 14 days on a numerical rating scale (NRS). PAF of the resting state electroencephalography (5 min) was assessed before and one day after the five rTMS treatment days.

The pre-registered analysis revealed no significant changes in PAF following five consecutive days of active (from 9.90±0.39 Hz to 9.95±0.38 Hz) or sham (from 9.86±0.44 Hz to 9.81±0.35 Hz) rTMS, suggesting that the impact of rTMS on NGF-induced pain is independent of PAF modulation. However, exploratory analysis indicated an association between a larger absolute difference in baseline PAF to 10 Hz (i.e. the rTMS frequency) and higher NRS pain ratings at Day 5 in participants receiving active rTMS. This suggests rTMS is more efficient when delivered close to individual PAF or for those with PAF around 10 Hz, necessitating further exploration of PAF’s role in rTMS-induced pain relief.

## 1 Introduction

Chronic pain imposes a significant burden on individuals, health care systems, and society, with pharmacological treatment demonstrating limited success (Cohen et al., 2021; Mann et al., 2013; Vos et al., 2012; Wewege et al., 2023). Repetitive transcranial magnetic stimulation (rTMS) has emerged as a promising adjunct treatment (Fernandes et al., 2022, 2022; Lefaucheur et al., 2020, 2014; Leung et al., 2009; Polanía et al., 2018). Using rTMS over the primary motor cortex (M1) or dorsolateral prefrontal cortex (DLPFC) produces sustained alleviation of pain in both chronic pain patients (Attal et al., 2021; Bursali et al., 2021; Mhalla et al., 2011; Passard et al., 2007) and healthy individuals undergoing experimentally-induced pain (Cavaleri et al., 2019; Ciampi De Andrade et al., 2014; De Martino et al., 2019; Nahmias et al., 2009; Seminowicz et al., 2018; Taylor et al., 2012). However, in a recent trial of 149 chronic pain patients, only 20–30% of patients described their pain as “much” or “very much” improved (Attal et al., 2021), suggesting that a substantial proportion of patients do not respond to rTMS treatment (Fernandes et al., 2022). Increased mechanistic understanding of how rTMS works for pain could allow appropriate selection of patients for treatment and enable individualisation of stimulation parameters for broader patient efficacy (Ciampi de Andrade and García-Larrea, 2023), however the mechanisms through which rTMS influences pain remain elusive (Fernandes et al., 2022). Given that the magnetic pulse of TMS induces electrical currents in the cortex that then influence the electrical activity of neural populations (Herrmann et al., 2016; Ridding and Rothwell, 2007), a potential factor to investigate in relation to the analgesic mechanisms of rTMS is peak alpha frequency (PAF), a neuronal oscillatory feature measured by electroencephalography (EEG).

Alpha oscillations (8–12 Hz) are the most visible feature of human brain activity recorded using EEG or magnetoencephalography (Van Diepen et al., 2019), with PAF being the specific frequency with maximal power within this alpha range. PAF increases during childhood, remains relatively stable with low intra-individual and high inter-individual variability during adolescence/adulthood (Bazanova and Vernon, 2014; Haegens et al., 2014), before decreasing during older age (Aurlien et al., 2004; Chiang et al., 2011; Lindsley, 1939). Similar to heart rate, intra-individual variation may indicate state-dependent adaptations or modifications, while inter-individual variation could represent underlying biological differences or variations in brain connectivity (Mierau et al., 2017). Inter-individual PAF variability has been linked to individual differences in visual and sensory processing (Bornkessel et al., 2004; Cecere et al., 2015; Doppelmayr et al., 2005; Jaušovec and Jaušovec, 2000), as well as pain sensitivity in both clinical and experimental settings (Chowdhury et al., 2025; De Martino et al., 2021; Fauchon et al., 2022; Furman et al., 2020, 2019, 2018; Lim et al., 2016; Mazaheri et al., 2022; Millard et al., 2022b; Nir et al., 2010; Sarnthein et al., 2006; Sufianov et al., 2014; Vries et al., 2013). Specifically, slower PAF is associated with chronic pain presence and duration (Fauchon et al., 2022; Lim et al., 2016; Sarnthein et al., 2006; Sufianov et al., 2014; Vries et al., 2013), as well as higher pain intensity ratings during prolonged experimental pain (Furman et al., 2020, 2019, 2018) and post-operative pain (Millard et al., 2022b).

Alongside PAF, studies have shown reduced alpha power during tonic or phasic experimental pain in healthy participants without chronic pain (Furman et al., 2021; Peng et al., 2014; Valentini et al., 2022) and, although not always consistent (Zebhauser et al., 2023), there is evidence for increased alpha power in chronic pain populations compared to healthy controls (Fauchon et al., 2022). Altered alpha power and frequency in those with chronic pain has been attributed the altered thalamocortical network excitability (Ahn et al., 2019; Fauchon et al., 2022; Peng et al., 2014). Therefore, as alpha oscillations may reflect or be involved in pain processing mechanisms (Mazaheri et al., 2022; Ploner et al., 2017), and rTMS has been shown to increase PAF (Anderson et al., 2007; Okamura et al., 2001), possibly by rhythmically influencing the electrical activity of cortical neural populations that give rise to oscillations (Herrmann et al., 2013; Polanía et al., 2018; Thut et al., 2011b; Vogeti et al., 2022), then facilitation of PAF could relate to the analgesic mechanism through which rTMS acts. However, it is currently unknown whether changes in PAF occur alongside the pain alleviating effects of rTMS.

This investigation could be conducted in two ways: 1) by taking an existing known effect of an intervention on pain and assessing whether there are concurrent effects on PAF, or 2) by taking an existing known effect on PAF and assessing whether there are concurrent effects on pain. Both approaches are valuable, complementary angles for investigating evidence for and use of the PAF–pain relationship. Examples of the latter approach include the use of nicotine to modulate PAF (Millard et al., 2023), where it is suggested that more direct PAF modulation methods, such as NIBS, may provide further insight. However as little research has been conducted on effective ways to modulate PAF using NIBS (Millard et al., 2024), the former approach was used here.

In this pre-registered secondary analysis (https://osf.io/mcjna/), we investigated whether PAF modulation occurs alongside established rTMS-induced pain modulation (De Martino et al., 2019; Seminowicz et al., 2018) in a randomised, single-blind, sham-controlled study. Healthy participants experienced experimentally induced sustained lateral epicondylalgia pain in the right extensor carpi radialis brevis (ECRB) muscle and underwent five days of high-frequency rTMS to the DLPFC, with resting state PAF measurements taken before (Day 0) and one day after (Day 5) the last rTMS session. Prior investigation using this dataset revealed that active rTMS reduced intensity ratings of the experimentally induced pain compared to sham rTMS (De Martino et al., 2019; Seminowicz et al., 2018). Here, it is hypothesised that active rTMS would 1) increase PAF, 2) increase power in the fast alpha band, and 3) decrease power in the slow alpha band relative to baseline compared to sham rTMS. We also included pre-registered exploratory analyses to assess whether potential changes in PAF were influenced by electrode location, pain level, and proximity of baseline PAF to the stimulation frequency of 10 Hz.

## 2 Methods

### 2.1 Participants

Thirty healthy right-handed female (n = 18) and male (n = 12) participants were recruited at Aalborg University with flyers and online adverts. Inclusion criteria: English speaking, age 21–50 years. Exclusion criteria: history of neurological, mental, or musculoskeletal illness; drug addiction (e.g., use of opioids, cannabis, or other drugs); known pregnancy; previous experience with rTMS; contraindications to rTMS based on the Transcranial Magnetic Stimulation Adult Safety Screen (Rossi et al., 2009). A physical examination was conducted prior to any experimental procedures to confirm the absence of tenderness to palpation of the ECRB muscle and the presence of full pain-free range of wrist and elbow motion. The original study was approved by local Ethics Committee (N-20170041), registered at ClinicalTrials.gov (NCT03263884), and was performed in accordance with the Helsinki Declaration. Informed written consent was obtained from each participant before study commencement.

This is a secondary analysis of data presented by Seminowicz et al. (2018) and De Martino et al. (2019), for which the EEG data is presented here. Previous outcomes of this dataset are published elsewhere, including effects of rTMS on pain, muscle soreness, painful area, disability, cognitive task performance (Seminowicz et al., 2018), pressure pain thresholds (PPTs), wrist extension force, sensory evoked potentials (SEPs), motor evoked potentials (MEPs), and coticospinal excitability measured by map volume (De Martino et al., 2019).

As described previously (Seminowicz et al., 2018), the sample included 30 healthy, rTMS naïve adults (mean age = 26.5 *±* 4.7 years), with 15 participants in each of the active (age = 26.9 *±* 3.8 years) and sham (age = 26.0 *±* 5.5 years) rTMS groups. Groups were matched by sex at birth (9 females and 6 males in each group). The intensity used for treatment was set at Day 0 and kept constant for the entire treatment, the rTMS intensity was 60.0 *±* 13.5% for the sham-rTMS group and 60.1 *±* 0.7% for the active-rTMS group. See Table 1 within Seminowicz et al. (2018) for full grouped participant information.

**Table 1:**
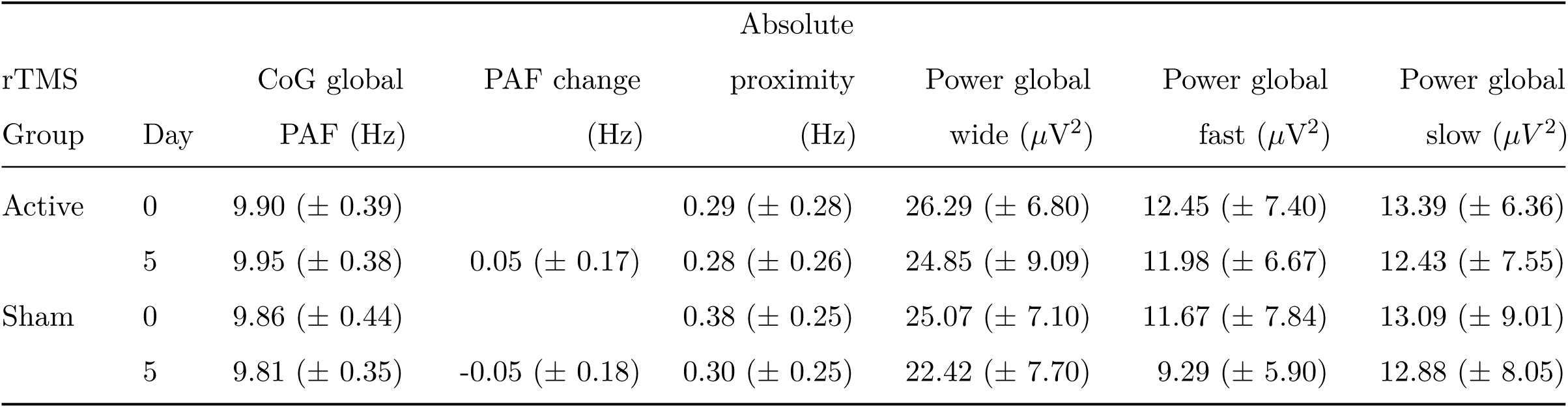
Mean *±* standard deviation (SD) peak alpha frequency (PAF), calculated with wide 8–12 Hz window and the centre of gravity (CoG) method. Power measured in *µV* ^2^ for wide band alpha (8–12 Hz) as well as in fast (10–11.8 Hz) and slow (8–9.8 Hz) alpha bands.

**Table 2:** Linear mixed-effects model results for the effect of Group (active vs. sham repetitive transcranial magnetic stimulation [rTMS]), Day (Day 5 vs. Day 0), and muscle soreness ratings on global 8–12 Hz peak alpha frequency (PAF). Estimates (*b−*coefficients), standard error (SE), 95% confidence intervals (CI), *t*-values, and *p*-values.

| Parameter | *b*-coefficients | SE | 95% CI | *t*-value | *p*-value |
| --- | --- | --- | --- | --- | --- |
| **NRS pain ratings** |  |  |  |  |  |
| *Intercept* | 9.425 | 0.250 | [8.956, 9.895] | 37.720 | <0.001 |
| Group (Active vs. Sham rTMS) | 0.374 | 0.327 | [-0.240, 0.987] | 1.145 | 0.262 |
| Day 5 (compared to baseline) | 0.117 | 0.110 | [-0.091, 0.325] | 1.060 | 0.299 |
| NRS (pain intensity on Day 5) | 0.117 | 0.061 | [0.001, 0.232] | 1.903 | 0.067 |
| Group * Day 5 | -0.119 | 0.144 | [-0.391, 0.153] | -0.823 | 0.418 |
| Group * NRS | -0.064 | 0.114 | [-0.277, 0.150] | -0.560 | 0.580 |
| Day 5 * NRS | -0.045 | 0.027 | [-0.096, 0.006] | -1.657 | 0.110 |
| Group * Day 5 * NRS | 0.069 | 0.050 | [-0.025, 0.164] | 1.379 | 0.180 |
| **Muscle soreness** |  |  |  |  |  |
| *Intercept* | 7.942 | 0.745 | [6.541, 9.342] | 10.654 | <0.001 |
| Group (Active vs. Sham rTMS) | 1.787 | 0.782 | [0.318, 3.256] | 2.285 | 0.030 |
| Day 5 (compared to baseline) | 0.409 | 0.354 | [-0.258, 1.077] | 1.156 | 0.258 |
| Soreness (muscle soreness on Day 5) | 0.457 | 0.176 | [0.126, 0.788] | 2.596 | 0.015 |
| Group * Day 5 | -0.357 | 0.372 | [-1.057, 0.344] | -0.959 | 0.346 |
| Group * Soreness | -0.390 | 0.195 | [-0.756, -0.025] | -2.004 | 0.054 |
| Day 5 * Soreness | -0.110 | 0.084 | [-0.267, 0.048] | -1.309 | 0.202 |
| Group * Day 5 * Soreness | 0.107 | 0.093 | [-0.068, 0.281] | 1.152 | 0.260 |

### 2.2 Experimental design

In this randomised sham-controlled, parallel design study, participants were screened for inclusion/exclusion over the phone and gave written informed consent prior to study commencement on Day 0. Participants attended six sessions over six consecutive days. An outline of the experiment timeline is displayed in Fig. 1. Resting state EEG was measured on Day 0, after which participants received five sessions of either active or sham rTMS on Days 0, 1, 2, 3, and 4. Resting state EEG was then measured again on Day 5. Additionally, participants received nerve growth factor (NGF) injections into their right ECRB muscle on Days 0 and 2 after the rTMS sessions to simulate sustained pain. Participants completed an online diary for 14 days to assess the pain in their elbow. Allocation to the active or sham-rTMS group was determined by a pre-set, randomly generated order. While participants remained blinded to allocation until study completion, the experimenter delivering rTMS was not. Each group comprised 15 participants, with nine females in each.

**Figure 1:**
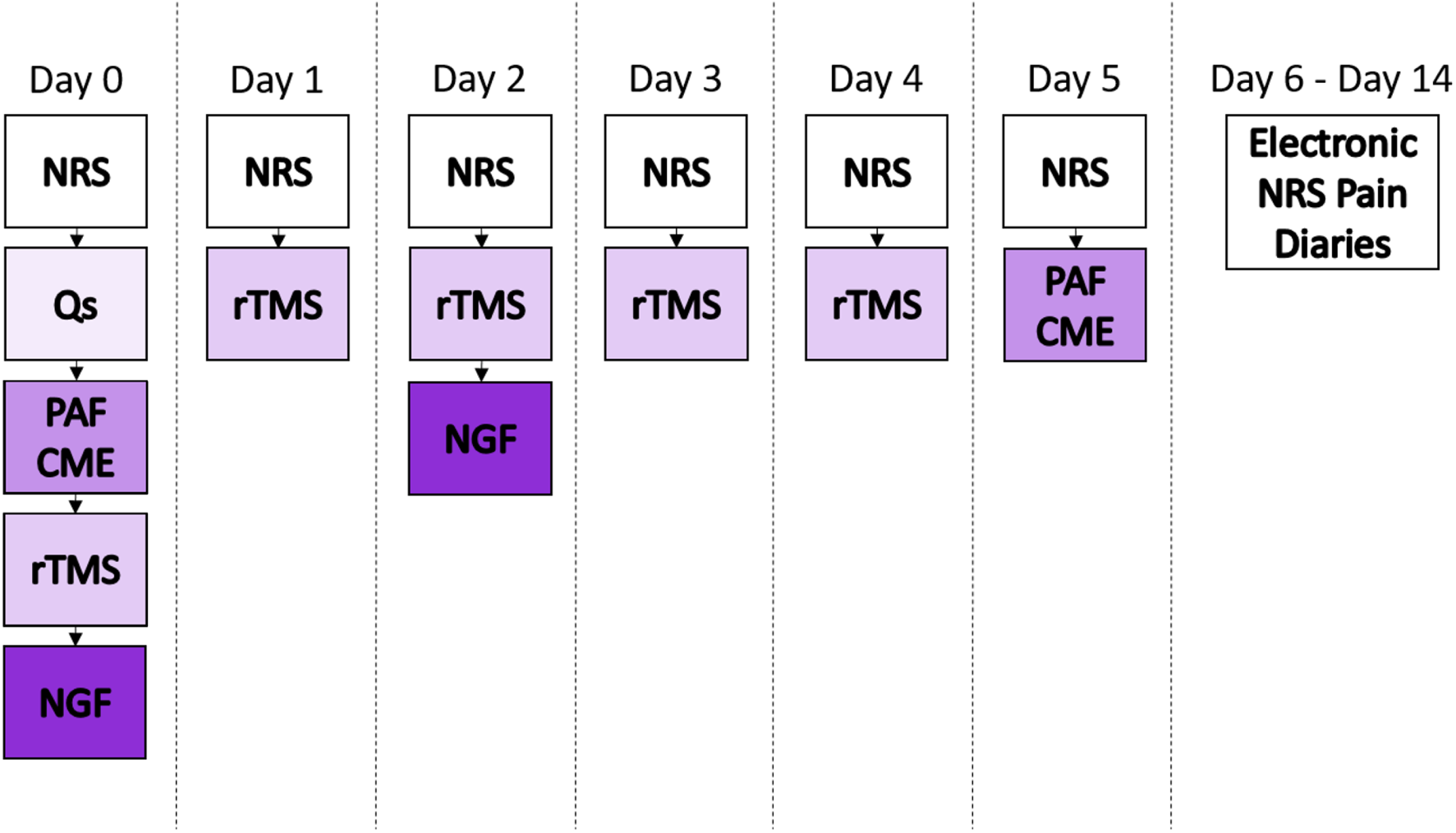
Timeline of experimental events. Participants reported their pain at the beginning of each experimental session on a numerical rating scale (NRS), and completed electronic pain diaries until Day 14. Questionnaires (Qs) were collected on Day 0. Electroencephalography (EEG) data were collected during eyes closed resting states on Day 0 and Day 5. Five consecutive days (Day 0–4) of repetitive transcranial magnetic stimulation (rTMS) were conducted. Nerve growth factor (NGF) was administered at the end of experimental sessions on Days 0 and 2.

Participants were screened for inclusion/exclusion over the phone and gave written informed consent prior to study commencement on Day 0. An outline of the experiment timeline is displayed in Fig. 1. Resting state EEG data was collected on Day 0 prior to rTMS and NGF injection, and again on Day 5 after participants had completed 5 sessions of stimulation to the DLPFC using active or sham rTMS (Days 0–4). The NGF injections took place after rTMS sessions on Days 0 and 2. Participants completed an online diary for 14 days to assess the pain in their elbow.

### 2.3 Data collection procedures

#### Questionnaires

Online pain diaries were completed on Days 0 to 14, in which pain severity and muscle soreness were assessed. An 11-point numerical rating scale (NRS) assessed pain severity from 0 = “no pain”, to 10 = “the worst pain imaginable”. A modified 7-point Likert scale assessed muscle soreness; 0 = “a complete absence of soreness,” 1 = “a light soreness in the muscle felt only when touched/vague ache,” 2 = “a moderate soreness felt only when touched/a slight persistent ache,” 3 = “a light muscle soreness when lifting or carrying objects,” 4 = “a light muscle soreness, stiffness, or weakness when moving the wrist without gripping an object,” 5 = “a moderate muscle soreness, stiffness, or weakness when moving the wrist,” 6 = “a severe muscle soreness, stiffness, or weakness that limits the ability to move.”

#### Electroencephalography

Surface EEG was collected with a 64 electrodes EEG cap (g.GAMMA cap2, Schiedlberg, Austria), in accordance with the international 10-20 system. Recordings were referenced online to an electrode placed on the right earlobe and the ground electrode was placed halfway between the eyebrows. A sampling rate of 2400 Hz was used, impedance was kept below 5 kΩ, and unfiltered EEG signals were amplified and digitised using a 50000x g.HIamp biosignal amplifier (g.tec-medical engineering GmbH, Schiedlberg, Austria). Once the EEG set-up was complete, participants were seated in a comfortable armchair in a quiet, semi-darkened room. A pillow around the neck was used to minimise the contraction of the neck muscles. The participants were instructed to keep their eyes closed during the continuous 5 minute EEG recording, remain still, and relax without falling asleep. Data were collected at the same time of day for each participant across the six sessions to minimise the influence of fluctuations in circadian rhythms.

#### Repetitive transcranial magnetic stimulation

rTMS was delivered in one session each day for 5 consecutive days (i.e. Day 0–4). Each stimulation session lasted 20 minutes and consisted of 80 trains of 5-second pulses applied at a frequency of 10 Hz, with a 10-second interval between trains, totalling 4000 pulses each day. This protocol ensured consistency with previous literature on rTMS and pain (Short et al., 2011; Taylor et al., 2012). The resting motor threshold (rMT) was determined for the right first dorsal interosseous muscle by visual inspection of the thumb movement and stimulation was applied at 110% of the rMT. A figure-of-eight shaped coil (70-mm Double Air Film Coil; Magstim Super Rapid2 Plus1, Magstim Co, Ltd, Dyfed, United Kingdom) was used and located using the BeamF3 algorithm (Beam et al., 2009; Mir-Moghtadaei et al., 2015; Seminowicz et al., 2018), a freely available tool (http://clinicalresearcher.org/software.htm). A sham coil (70-mm Double Air Film Sham Coil) of the same shape, size, and colour, which produced a similar sound to the active coil, was used for sham stimulation. The instructions to participants and procedures were identical for both the active and sham rTMS groups.

#### Experimentally induced pain

NGF was injected into the right ECRB muscle on Days 0 and 2 to induce muscle pain and hyperalgesia, modelling lateral epicondylalgia (Bergin et al., 2015). A pharmacy (Skanderborg Apotek, Skanderborg, Denmark) prepared sterile solutions of recombinant human NGF. After the site had been cleaned with alcohol, and guided by in-plane real-time ultrasound (SonoSite M-Turbo; FUJIFILM SonoSite, Inc), the NGF solution (5 *µ*g/0.5 mL) was injected into the ECRB muscle belly. During experimental sessions, participants were asked to rate their pain at rest as well as during maximal wrist extension, holding a weight (1.3 kg) with the forearm supported on a platform in pronation (contraction-state).

### 2.4 Data processing

#### EEG pre-processing

Pre-processing of de-identified EEG data was completed using custom code in MATLAB 2022b (v. 9.13.0.2126072, Update 3, The MathWorks, Inc., Natick, MA, USA) using EEGLAB 2022.1 (Delorme and Makeig, 2004) and FieldTrip (v.20210929) (Oost-enveld et al., 2011). The experimenter conducting pre-processing was blinded to participant groups. Using EEGLAB, for resting state EEG recordings on Days 0 and 5, data were downsampled to 500 Hz, filtered between 2 and 100 Hz using a linear finite impulse response (FIR) bandpass filter, and the 5 minute period of eyes closed resting state were visually inspected and overtly noisy channels with *≥* 8 signal deviations (e.g., popping, baseline drift, line noise) were removed. The average number of channels removed were 0.94 and 0.87 (range: 0–4) for the two resting states, respectively. Removed channels were interpolated based on the nearest neighbour method (Chowdhury et al., 2023). Data were re-referenced to the common average reference.

Using FieldTrip, data were segmented into 5 second epochs, with no overlap, visually inspected, and epochs containing any electrode, muscle, or motion artefacts – other than eye blinks or saccades – were removed. Principal component analysis using the *runica* method was applied to identify and remove components reflecting eye blinks and/or saccades. The average number of components removed per resting state was 1.05 and 1.2 (range: 0–2), respectively.

From a maximum of 60 epochs (i.e. 5 minutes), the average number of remaining epochs was 39.00 and 40.10 (range: 36–51) for each resting state, respectively. Therefore, no resting state contained less than three minutes of data, thus producing more reliable PAF values, for which at least two minutes is advised (Chowdhury et al., 2023).

#### Quantification of EEG outcomes

In this analysis, primary outcomes included PAF, as well as power in two distinct alpha oscillation bands: slow (i.e. < 10 Hz) and fast (i.e. > 10 Hz) alpha oscillations. These three primary outcomes were estimated for the two eyes closed resting states (i.e. pre-rTMS and post-rTMS). All measures were calculated by averaging across all sensors to obtain global measures.

FieldTrip was used to calculate the power spectral density in the range of 2–40 Hz for each epoch in 0.2 Hz bins. A Hanning taper was applied to the data prior to calculating spectra to reduce any edge artefacts (Mazaheri et al., 2014). Band power was quantified as the z-scored total spectral density within the band, measured in *µ*V^2^. The use of split power bands stemmed from the understanding that PAF may reflect the balance between slow and fast alpha power, with shifts in PAF potentially arising from changes in either or both power bands (Furman et al., 2021). Note that we defined the slow alpha band as frequencies between and including 8–9.8 Hz and the fast alpha band as frequencies between and including 10–11.8 Hz. This choice was pre-registered and made due to the 0.2 Hz frequency resolution, to avoid overlap at 10 Hz, and to ensure the same number of frequency bins in each band, therefore power at 12 Hz was omitted.

The frequency range used to calculate PAF varies in the PAF–pain literature. The narrower 9–11 Hz band reduces the impact of 1/f noise (i.e. trend of higher power at lower frequencies) on the PAF estimations (Furman et al., 2018; Mazaheri et al., 2014); however the narrow band may not produce representative PAF values for those with peaks near the lower or upper limits. Therefore, the wider 8–12 Hz band was used as the primary band of interest.

Similar to previous (Chowdhury et al., 2023; Furman et al., 2020, 2019, 2018; Millard et al., 2022b), the PAF for each power spectral density was estimated using the centre of gravity (CoG) method (Brötzner et al., 2014; Jann et al., 2012, 2010; Klimesch, 1999; Klimesch et al., 1993):

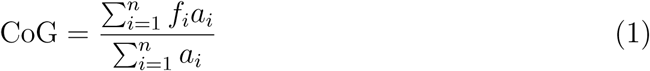

where *f_i_* is the *i*^th^ frequency bin, *n* is the number of frequency bins within the frequency window (8–12Hz), and *a_i_* is the amplitude of the frequency for *f_i_*. This equation states that PAF is the weighted sum of spectral estimates (Klimesch, 1999).

The peak detection method (i.e., identifying the frequency with maximum power within the alpha range) was calculated in addition to the CoG method as an exploratory validation check. Additionally, a sensorimotor region of interest (ROI; i.e. Cz, C1, C2, C3, C4) was also analysed due to its frequent use in PAF–pain research (Chowdhury et al., 2025, 2023; Furman et al., 2020, 2019, 2018).

### 2.5 Statistical analysis

Statistical analysis was conducted in RStudio (RStudio Team, 2022.4.2.1). Mean and standard deviations (SD) for primary outcomes were calculated for each group (i.e. active and sham rTMS groups). Any data point three SDs above or below the mean of the respective group was removed. Data were analysed using mixed model analysis of variances (ANOVAs) to assess any changes in each measure over time given the intervention group. Assumptions were assessed using Shapiro-Wilk tests, Q-Q plots, and histograms for normality. All statistical analyses report associated *F* statistics, *p*-values (*α* = .05), standard errors, as well as corresponding 95% confidence intervals unless stated.

A mixed model ANOVA assessed whether any increase in PAF speed for active rTMS was larger than for sham rTMS: with PAF as the dependent variable, Day (2 levels: Day 0/Day 5) as the within participant variable and Group (2 levels: active/sham rTMS) as the between participant variable. The key effect examined was the interaction between group and day, with planned pairwise comparisons of group and time point. To assess fast alpha power increases and slow alpha power decreases in response to active rTMS, separate mixed model ANOVAs were conducted with global power in the fast or slow alpha band as the dependent variables.

A planned exploratory analysis was to assess the three primary hypotheses whilst additionally controlling for pain levels produced by experimentally induced muscle pain. Linear mixed-effects models were used to investigate the effects of group, day, and pain intensity at Day 5 on the global PAF, with participant ID included as a random effect. The model formula was PAF ∼ Group * Day * pain + (1 | Participant_ID), the reference group was set to sham rTMS, and the analysis was conducted twice: once using NRS ratings as the pain variable and again using muscle soreness ratings as the pain variable. The analysis was conducted using the lmer function from the lme4 package in R.

In a second planned exploratory analysis, the difference in baseline PAF from 10 Hz (absolute proximity) was measured in Hz and was calculated for each participant for Day 0 and 5 (i.e. pre-rTMS and post-rTMS). Correlations were conducted to assess the relationship between PAF distance from 10 Hz at baseline and a) change in PAF, as well as b) NRS on Day 5. Day 5 was chosen because a previous study assessing pain in the ECRB muscle after NGF injection showed an average peak NRS pain on Day 5 (De Martino et al., 2019; Furman et al., 2019).

Lastly, as a global PAF measure including all electrodes is not able to capture spatial variations in PAF changes, we conducted a data-driven exploratory non-parametric cluster-based permutation analysis (CBPA) (Maris and Oostenveld, 2007; Pernet et al., 2015). This approach allows for the detection of spatially contiguous effects by assessing statistical differences in PAF pre-/post-rTMS across the scalp, akin to methods used by Furman et al. (2021) and Millard et al. (2023) to assess changes in PAF. Change in PAF from baseline to post-rTMS was assessed for the sham and active rTMS groups separately.

The CBPA was conducted as follows, separately for the sham and active rTMS groups:

1. Initial single-electrode tests: A paired t-test was performed at each electrode to compare pre-/post-rTMS.
2. Thresholding: Electrodes showing differences at *p < .*01 on the statistical test were considered for clustering.
3. Cluster formation: If two or more spatially adjacent electrodes passed the threshold they were grouped into a cluster. Each electrode within a cluster had to also be adjacent to at least one other electrode from the the cluster.
4. Cluster-level statistic: The test statistic values (i.e. t-values) from all electrodes within each cluster were then summed to give an overall cluster-level test value for each separate cluster.
5. Permutation testing: To establish statistical significance, a null distribution was generated by randomly shuffling dependent variable values 1000 times, recalculating the cluster-level test statistic for each permutation.
6. Significance determination: If the observed cluster-level statistic was greater than the null distribution 99th percentile, that cluster was deemed statistically significant.

## 3 Results

As previously described, rTMS to the left DLPFC was associated with reduced pain intensity, painful area, and muscle soreness compared to the sham group (Seminowicz et al., 2018), as well as higher PPTs and muscle pain scores in the active compared to sham group (De Martino et al., 2019). The mean and SD values for pain outcomes are reported in Supplementary Table S1 made available via Seminowicz et al (2018) (http://links.lww.com/PAIN/A636). The present secondary analysis assessed whether there were concurrent effects of five days of left DLPFC rTMS on resting state EEG alongside the effects on pain observed in this dataset.

### 3.1 Change in peak alpha frequency

PAF and power values are shown in Table 1, with individual and average spectra by group in Fig. 2, and overlaid averaged spectral and topographical plots by Day and group displayed in Fig. 3. Following CONSORT recommendations (Moher et al., 2012; Pijls, 2022), no statistical tests of baseline equivalence were performed.

**Figure 2:**
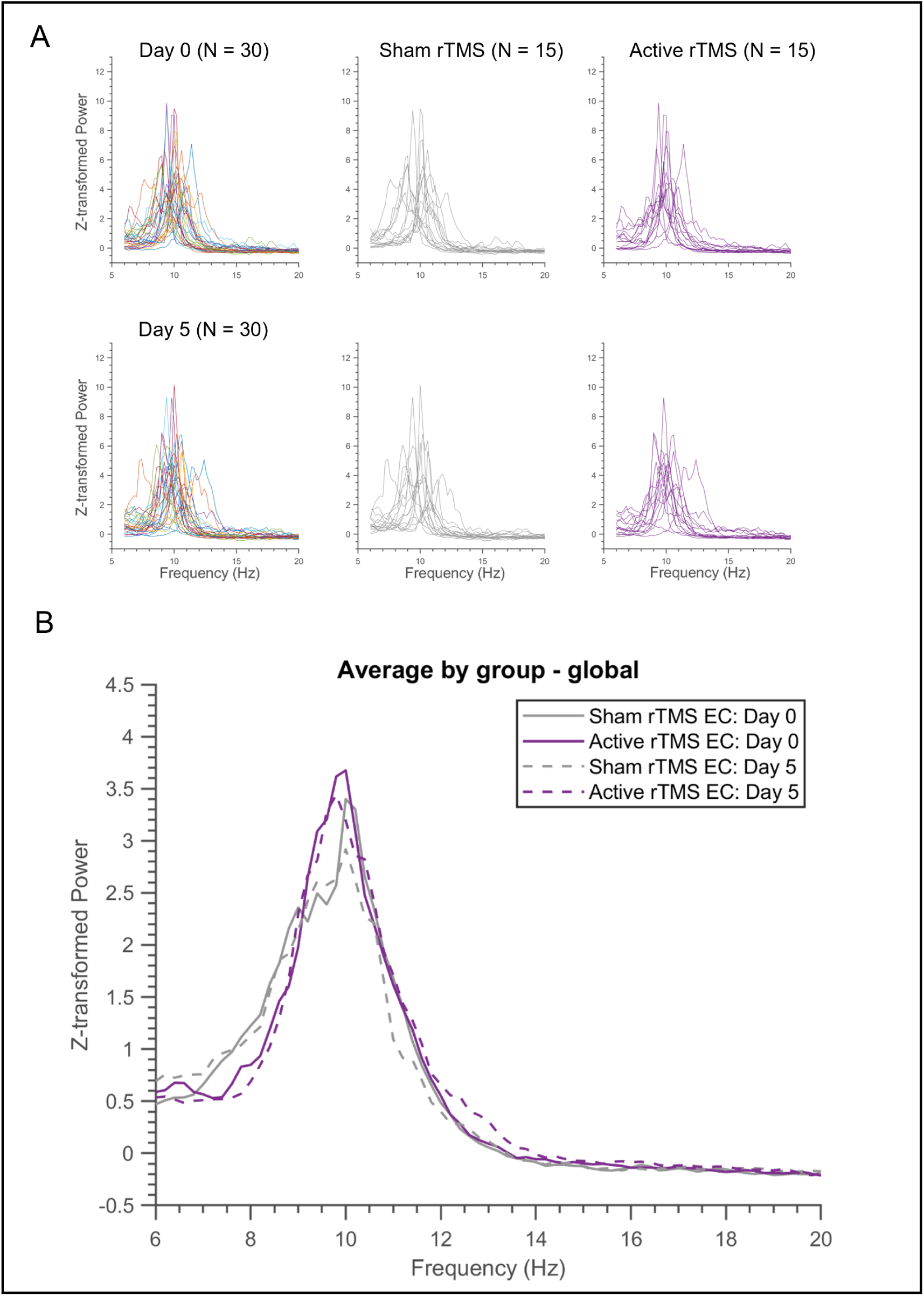
Individual and averaged Z-transformed global power spectra of electroen-cephalography (EEG) recordings on Day 0 and Day 5 across groups. **A)** The top panel in A represents EEG spectral power on Day 0 (before repetitive transcranial magnetic stimulation [rTMS] and induction of sustained muscle pain via nerve growth factor [NGF]), while the bottom panel in A represents EEG spectral power on Day 5 (after five consecutive days of rTMS and sustained muscle pain). The first column in A shows individual spectra for all participants (N = 30). The second and third columns display individual spectra for participants receiving sham rTMS (n = 15, grey) and active rTMS (n = 15, purple), respectively. **B)** Group-averaged spectra pre-(solid line) and post-rTMS (dashed line) for sham (grey) and active (purple) rTMS.

**Figure 3:**
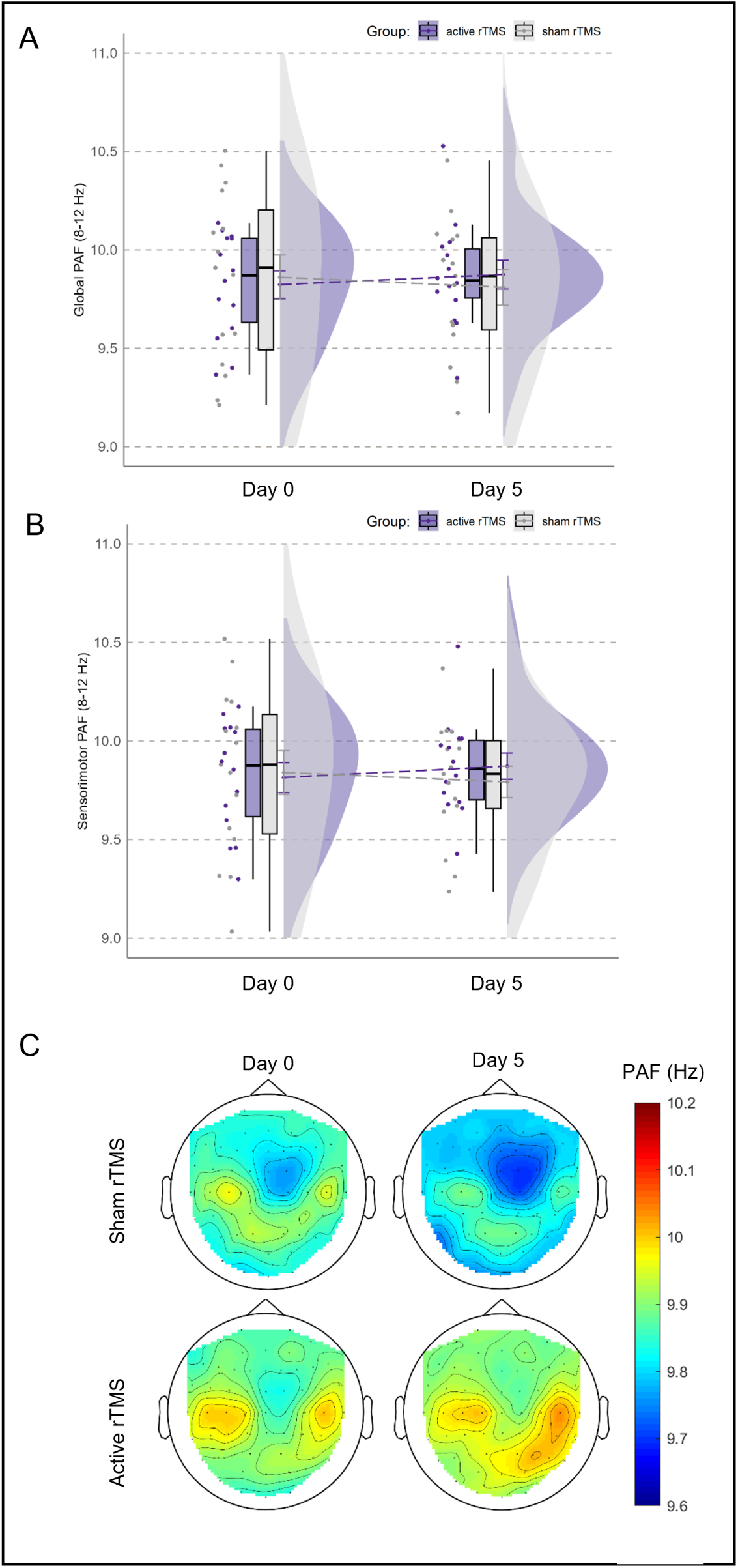
**A)** Raincloud plots of mean global peak alpha frequency (PAF) values before and after either sham (grey) or active (purple) rTMS. **B)** Raincloud plots of mean sensorimotor PAF values (i.e. Cz, C1, C2, C3, C4) before and after either sham (grey) or active (purple) rTMS. **C)** Topographical plots of center of gravity (CoG) PAF calculated for each electrode for Day 0 (baseline) and Day 5 by sham (top panel) and active rTMS groups (bottom panel).

One participant was excluded from the analysis of active-rTMS effects on PAF because their PAF value was >3 SD above the mean. PAF data within Group and Day were normally distributed (*W* = 0.91 *−* 0.98*, p > .*16). For the effects of rTMS on PAF, there were no significant main effects of Group (*F* [1, 27] = 0.01*, p* = .91, 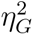 *<* 0.01) or Day (*F* [1, 27] *<* 0.01*, p* = .99, 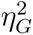 *<* 0.01) and no Group by Day interaction (*F* [1, 27] = 2.39*, p* = .13, 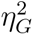 = 0.01; Fig. 3).

The exploratory linear regression model controlling for the potential effect of Day 5 NRS pain ratings on PAF at Day 5 (*R*^2^ = 0.77) demonstrated a significant estimated effect of baseline PAF (*b* = 0.76*, p < .*001), but the estimated effects of Group (*b* = *−*0.075*, p* = .32) and Day 5 NRS pain ratings were non-significant (*b* = *−*0.00092*, p* = .023).

#### Comparison of PAF calculation methods and location

PAF estimation using the CoG method was highly correlated with PAF calculated using the classical peak detection method across all conditions (global and sensorimotor PAF on days 0 and 5; all *r >* 0.97, all *p* = 2.2 × 10*^−^*^16^). The primary findings regarding lack of an effect of sham versus active rTMS on PAF modulation remained unchanged.

Specifically, there were no significant main effects of Group (*F* [1, 27] = 0.036*, p* = .85, 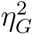 *<* 0.01) or Day (*F* [1, 27] = 0.009*, p* = .93, 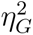 *<* 0.01) and no Group by Day interaction (*F* [1, 27] = 1.89*, p* = .18, 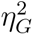 = 0.01) for global PAF with the peak picking method. There were no significant main effects of Group (*F* [1, 27] = 0.056*, p* = .82, 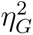 *<* 0.01) or Day (*F* [1, 27] = 0.018*, p* = .89, 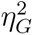 *<* 0.01) and no Group by Day interaction (*F* [1, 27] = 2.19*, p* = .15, 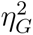 = 0.01) for sensorimotor PAF with CoG method. Lastly, there were no significant main effects of Group (*F* [1, 27] = 0.012*, p* = .91, 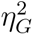 *<* 0.01) or Day (*F* [1, 27] = 0.044*, p* = .84, 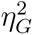 *<* 0.01) and no Group by Day interaction (*F* [1, 27] = 2.97*, p* = .097, 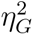 = 0.01) for sensorimotor PAF with the peak picking method.

A cluster-based permutation analysis was conducted to indicate whether there were more fine-grained locations for PAF change, but this also suggested no significant clusters of electrodes with PAF changes in the active or sham rTMS groups.

### 3.2 Change in fast and slow alpha power

Wide band power data within Group and Day were normally distributed (*W* = 0.88 *−* 0.97*, p > .*053). For the effects of rTMS on power across the whole alpha band (8–12 Hz), there were no significant main effects of Group (*F* [1, 28] = 0.54*, p* = .47, 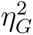 = 0.015) or Day (*F* [1, 28] = 0.14*, p* = .14, 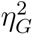 = 0.018) and no Group by Day interaction (*F* [1, 28] = 0.20*, p* = .66, 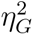 *<* 0.01).

Slow alpha band power data within Group and Day were normally distributed (*W* = 0.91 *−* 0.95*, p > .*13). For the effects of rTMS on slow alpha power, there were no significant main effects of rTMS group (*F* [1, 28] *<* 0.01*, p* = .98, 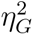 *<* 0.01) or Day (*F* [1, 28] = 0.80*, p* = .38, 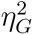 *<* 0.01) and no interaction (*F* [1, 28] = 0.33*, p* = .57, 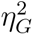 *<* 0.01; Fig. 4). The exploratory linear regression model for the effect of pain level on slow alpha power (*R*^2^ = 0.90) demonstrated a significant estimated effect of baseline slow alpha on Day 5 slow alpha power (*b* = 0.76*, p < .*001), but the estimated effects of Group (*b* = 0.30*, p* = .85) and Day 5 NRS pain ratings were non-significant (*b* = 0.24*, p* = .64).

**Figure 4:**
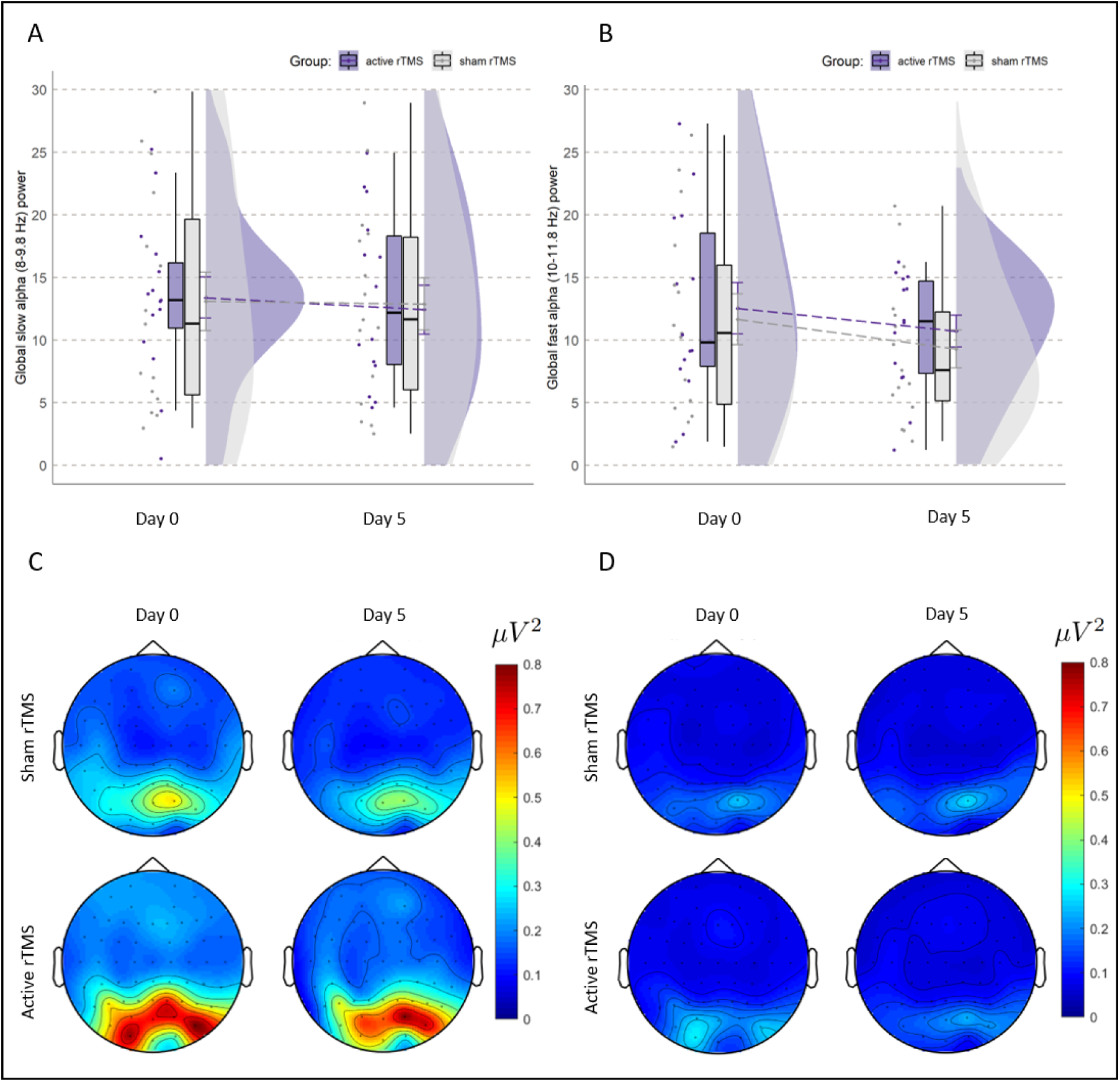
**A)** Raincloud plots of mean global power values in the slow alpha band (8–9.8 Hz), before and after either active (in purple) or sham (in grey) rTMS. **B)** Raincloud plots of mean global power values in the fast alpha band (10–11.8 Hz), before and after either active (in purple) or sham (in grey) rTMS. **C)** Topographical plots of power calculated for each electrode across the slow alpha band (8–9.8 Hz) are shown for the Day 0 (baseline) and Day 5. **D)** Topographical plots of power calculated for each electrode across the fast alpha band (10–11.8 Hz) are shown for Day 0 (baseline) and Day 5.

One participant was excluded from the analysis of rTMS effects on fast alpha power because their power value was >3 SD above the mean. After removing this participant, fast alpha band power data within Group and Day were normally distributed (*W* = 0.91 *−* 0.94*, p > .*17). For the effects of rTMS on fast alpha power, there was a main effect of Day (*F* [1, 27] = 5.62*, p* = .03, 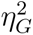 = 0.03) with no main effect of Group (*F* [1, 27] = 0.25*, p* = .62, 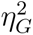 = 0.01) and no Group by Day interaction (*F* [1, 27] = 0.10*, p* = .76, 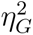 *<* 0.01; Fig. 4). The exploratory linear regression model for the influence of pain level on fast alpha power after rTMS (*R*^2^ = 0.64) demonstrated a significant estimated effect of baseline fast alpha on Day 5 fast alpha power (*b* = 0.55*, p < .*001), but the estimated effects of Group (*b* = *−*0.95*, p* = .54) and Day 5 NRS pain ratings were non-significant (*b* = 0.0015*, p > .*99).

### 3.3 Exploratory analysis of absolute proximity

One participant was removed from the active rTMS group due to having a proximity value > 3 SD above the mean. There was a correlation between absolute proximity (i.e. the difference between baseline PAF and 10 Hz) at Day 0 (baseline) and NRS pain on Day 5 in the active rTMS group (*r* = 0.55*, p* = .042, 95% CI: [0.025, 0.84]), but not the sham group (*r* = *−*0.12*, p* = .67, 95% CI: [-0.60, 0.42]; Fig. 5). This suggests that participants in the active rTMS group whose baseline PAF was closer to 10 Hz reported less pain on Day 5. Both the active and the sham groups had nine participants with global PAF (8–12 Hz) values below 10 Hz, and therefore had five and six participants with PAF values above 10 Hz, respectively.

**Figure 5:**
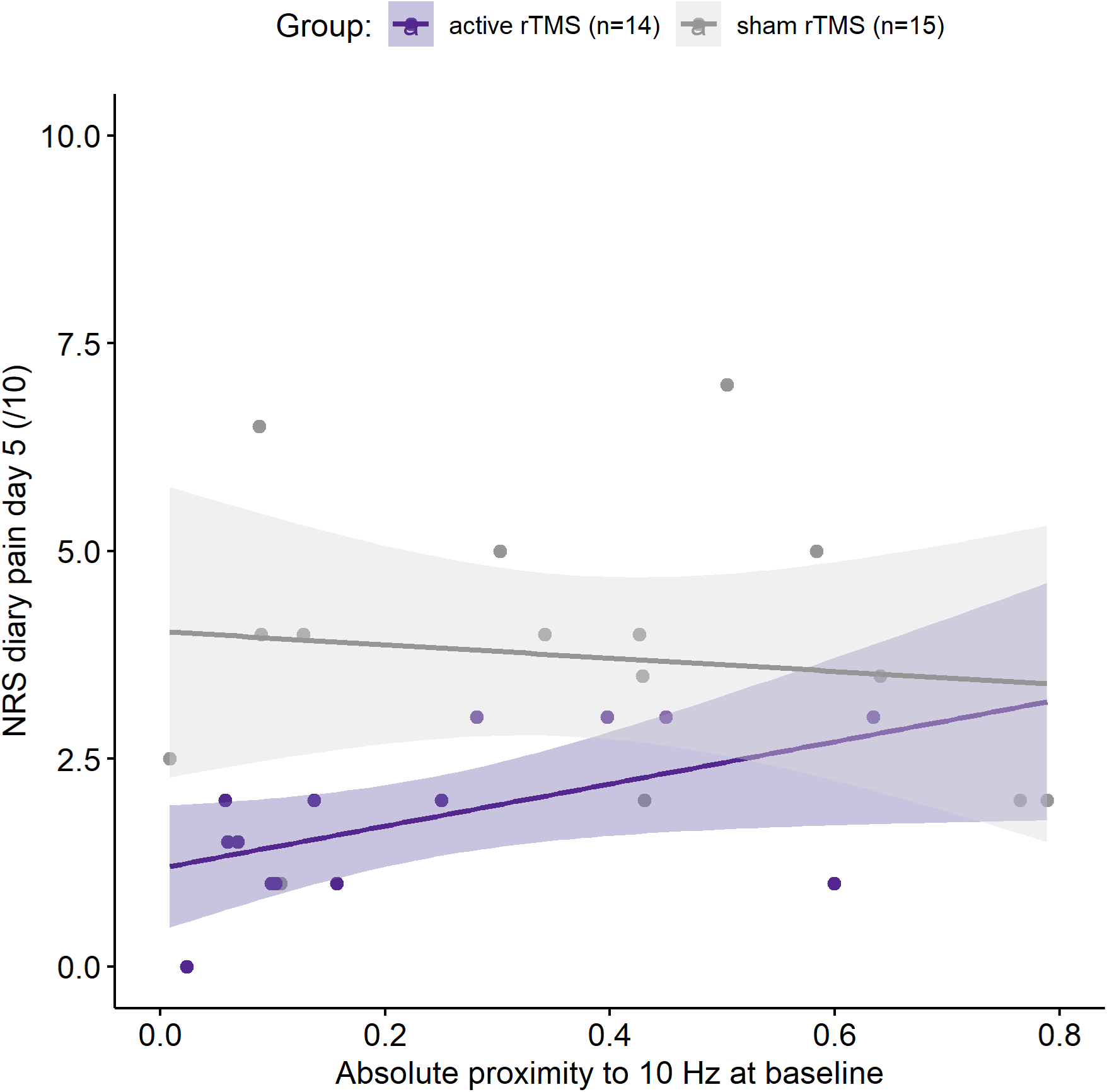
Relation between distance of baseline wide band (8–12 Hz) global PAF to the stimulation frequency of 10 Hz (i.e. absolute proximity) and NRS pain on Day 5. Separated into sham (*r* = *−*0.12*, p* = .67, 95% CI: [-0.60, 0.42]) and active (*r* = 0.55*, p* = .042, 95% CI: [0.025, 0.84]) rTMS groups. Regression lines and shaded 95% confidence intervals.

### 3.4 Correlations between PAF and pain

Baseline PAF was not correlated with Day 5 NRS pain ratings for the whole sample (N=29, *rs* = 0.11*, p* = .59), or to muscle soreness on day 5 (N=29, *rs* = 0.23*, p* = .23). When stratified by stimulation groups (Fig. 6), there were also no relationships between baseline PAF and NRS ratings on Day 5 for the active rTMS group (n=14, *rs* = *−*0.41*, p* = .14) or the sham rTMS group (n=15, *rs* = 0.43*, p* = .11). However, although there was still no correlation for the active group between baseline PAF and muscle soreness on Day 5 (N=14, *rs* = 0.10*, p* = .73), there was a significant correlation for the sham group (N=15, *rs* = 0.56*, p* = .031).

**Figure 6:**
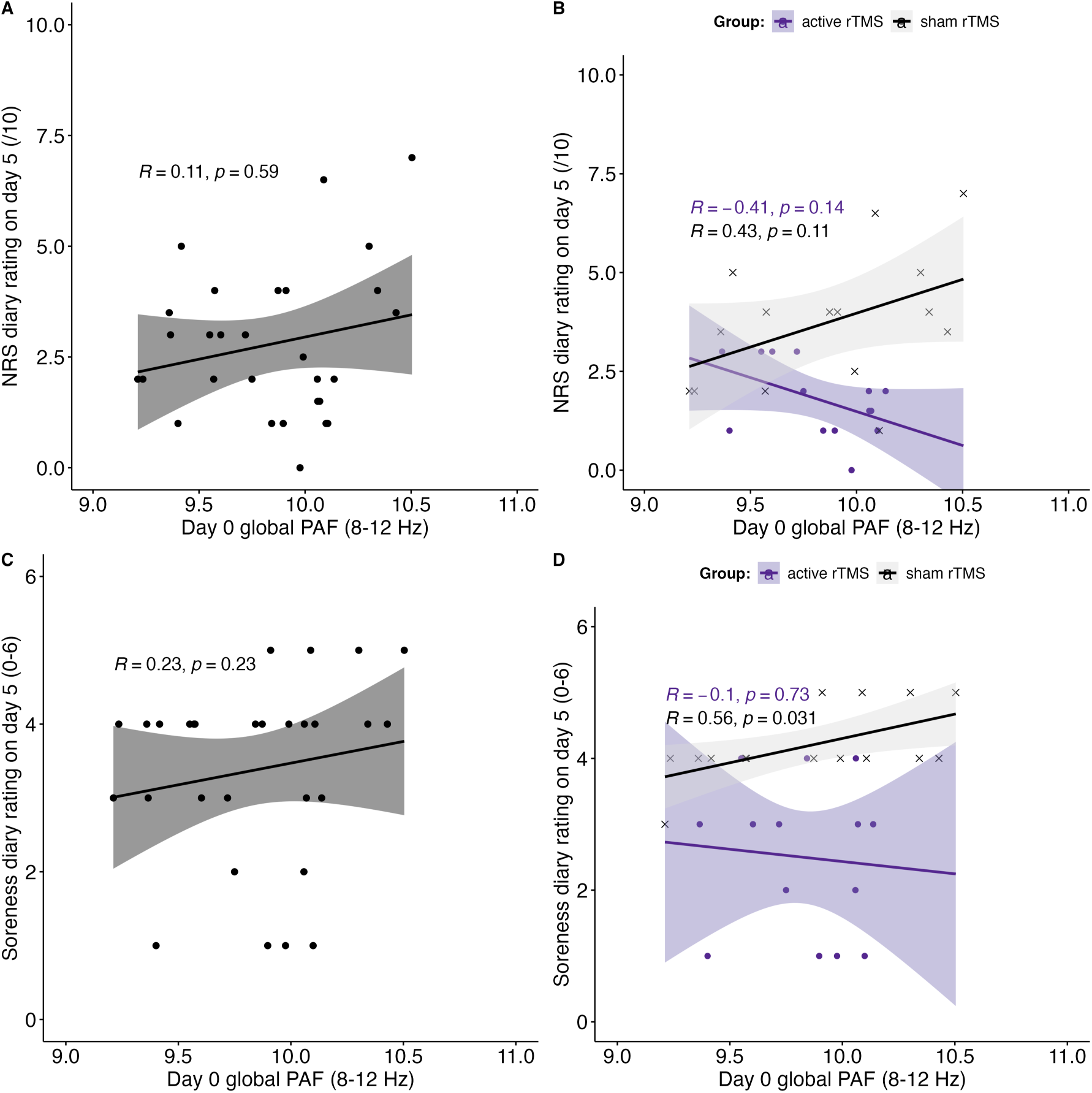
Baseline wide band (8–12 Hz) global peak alpha frequency (PAF) does not correlate with pain rated on a numerical rating scale (NRS) on Day 5, for **A)** the whole sample (N=30), or **B)** separately for the active (n=14) and sham (n=15) rTMS groups. Baseline PAF also does not correlate with muscle soreness on Day 5 for **C)** the whole sample (N=30). But when assessed **D)** separately for the active (n=14) and sham (n=15) rTMS groups, there is a positive correlation within the sham group, suggesting that higher baseline PAF is related to higher muscle soreness after sham rTMS. Regression lines and shaded 95% confidence intervals.

### 3.5 Exploratory analysis of effects of rTMS and pain intensity on PAF

Linear mixed-effects models were used to investigate the effects of group, day, and pain intensity at Day 5 on the global PAF, with participant ID included as a random effect. The model formula was PAF ∼ Group * Day * pain + (1 | Participant_ID), the reference group was set to sham rTMS, and the analysis was conducted twice: once using NRS ratings as the pain variable and again using muscle soreness ratings as the pain variable.

For the NRS pain rating model, as the intercept, the mean global PAF at baseline for the sham rTMS group was approximately 9.43 (95% CI: [9.032, 9.818], *t* = 44.91). The marginal *R*^2^ value indicates that 15.1% of the variance in global PAF is explained by the fixed effects, whereas the conditional *R*^2^ value suggests that 88.2% of the variance is explained when considering both fixed and random effects. The effect of active rTMS compared to sham rTMS on global PAF was positive and statistically significant (*b* = 0.63, 95% CI: [0.10, 1.16], *t* = 2.23, *p* = .034), suggesting that PAF was higher in the active rTMS group. Changes in global PAF from baseline to Day 5 were not statistically significant (*b* = 0.12, 95% CI: [-0.091, 0.33], *t* = 1.06, *p* = .30). The effect of pain intensity at Day 5 on PAF was significant (*b* = 0.12, 95% CI: [0.02, 0.21], *t* = 2.27, *p* = .031), indicating that higher pain intensity is associated with higher global PAF. The interaction between Group and NRS pain intensity was also significant (*b* = -0.25, 95% CI: [-0.45, -0.046], *t* = -2.298, *p* = .029). This significant negative interaction term suggests that the relationship between pain ratings and PAF differs between the sham and active rTMS groups. The main effect of NRS pain ratings suggests that there is a positive relation between pain ratings and PAF in the sham group, whereas in the active rTMS group, this relationship is less positive or possibly negative. This indicates that the active rTMS group does not follow the same pattern as the sham group and might even exhibit an inverse relationship between pain ratings and PAF. Further studies may be needed to explore potential nuanced effects. None of the other interactions between Group, Day, and NRS pain intensity were statistically significant.

For the muscle soreness model, as the intercept, the mean global PAF at baseline for the sham rTMS group was approximately 7.94 (95% CI: [6.76, 9.12], *t* = 12.61). The marginal *R*^2^ value indicates that 21.3% of the variance in global PAF is explained by the fixed effects, whereas the conditional *R*^2^ value suggests that 87.2% of the variance is explained when considering both fixed and random effects. The effect of active rTMS compared to sham rTMS on global PAF was again positive and statistically significant (*b* = 1.93, 95% CI: [0.69, 3.17], *t* = 2.91, *p* = .007). Changes in global PAF from baseline to Day 5 were not statistically significant (*b* = 0.41, 95% CI: [-0.27, 1.09], *t* = 1.14, *p* = .27). The effect of muscle soreness at Day 5 on PAF was significant (*b* = 0.46, 95% CI: [0.18, 0.74], *t* = 3.07, *p* = .005), suggesting that higher muscle soreness is associated with higher PAF. And similarly to the NRS pain model, the interaction between Group and muscle soreness was significant (*b* = -0.48, 95% CI: [-0.79, -0.17], *t* = -2.86, *p* = .008). None of the other interactions between Group, Day, and muscle soreness were statistically significant.

## 4 Discussion

In this pre-registered analysis, we probed the proposed PAF–pain relationship (Furman et al., 2020, 2019, 2018; Mazaheri et al., 2022; Millard et al., 2022b) by investigating whether modulation of EEG PAF or alpha power occurred alongside established DLPFC rTMS-induced reduction of sustained lateral epicondylalgia pain after NGF injection to the right ECRB muscle (Seminowicz et al., 2018). Contrary to predictions, we found that active rTMS did not significantly increase PAF from baseline compared to sham rTMS, despite effects on NGF-induced pain. In addition, there were no significant changes in wide band (8–12 Hz) alpha or slow (8–9.8 Hz) alpha power, but fast (10–11.8 Hz) alpha power decreased during pain irrespective of active of sham-rTMS. Therefore, changes in PAF speed and alpha power did not co-occur with rTMS-induced pain relief. However, our exploratory analysis found that the closer an individual’s baseline PAF was to 10 Hz, the greater the analgesic effect of active 10 Hz rTMS, and that the relationship between PAF and pain may have differed for the active and sham rTMS groups.

### 4.1 rTMS did not increase PAF despite influencing pain

Few studies have assessed state changes in PAF related to pain in a controlled experimental setting (Millard et al., 2023; Sato et al., 2021), with most research assessing concurrent changes coming from the chronic pain literature (Heitmann et al., 2022; Ngernyam et al., 2015; Parker et al., 2021; Sarnthein et al., 2006; Sato et al., 2017; Sufianov et al., 2014). In this study, we conducted a secondary analysis to examine the effects of 10 Hz rTMS over the DLPFC on PAF alongside the known effects on experimentally induced NGF-pain (De Martino et al., 2019; Seminowicz et al., 2018), which is the first investigation using rTMS and experimental pain to investigate the PAF–pain relationship. Contrary to expectations, no significant increases in global PAF were observed on the day following five consecutive days of rTMS, with no electrode clusters demonstrating substantial alterations. These findings, in concert with the previously published analgesic effects of active rTMS compared to sham rTMS in the same dataset (De Martino et al., 2019; Seminowicz et al., 2018), suggest that while PAF has shown promise in distinguishing high from low pain sensitive individuals (Furman et al., 2020, 2019, 2018; Mazaheri et al., 2022; Millard et al., 2022b), changes in experimental pain sensitivity using rTMS do not necessarily correspond with state changes in PAF.

Previous literature has demonstrated changes in PAF alongside changes in experimental pain using nicotine (Millard et al., 2023) and exercise alone, as well as exercise combined with transcranial direct current stimulation (tDCS) (Sato et al., 2021), but not with tDCS alone (Sato et al., 2021) or rTMS in the present study. To understand how different interventions have selective influence on pain and/or PAF, we need to consider the complex and distributed nature of pain processing. Nociceptive signals and pain are processed at multiple levels within the spinal cord and brain, making the pain system resilient to disruption (Coghill, 2020). This resilience is evidenced by the fact that pain can originate through various independent brain mechanisms (Coghill, 2020; Coghill et al., 1999; Dum et al., 2009; Finnerup et al., 2020; Noordenbos and Wall, 1976; Sugar and Bucy, 1951). PAF may represent one aspect of this system, explaining the mixed results regarding concurrent changes in PAF and experimental pain. By examining which interventions produce concurrent effects and which do not, we may identify the specific aspects of the pain system that PAF reflects. Given the present results, alongside the existing literature (Millard et al., 2023; Sato et al., 2021), we could suggest that broad interventions like exercise and nicotine target aspects of pain processing associated with PAF, whereas those with a narrower spatial resolution, such as rTMS, may act on mechanisms independent of PAF (see Fernandes et al. (2022) and Ciampi de Andrade and García-Larrea (2023) for recent reviews of mechanisms of rTMS for pain). Therefore, PAF likely reflects only a subset of the processes contributing to pain sensitivity, while other neurophysiological, cognitive, or psychological factors may also play significant roles. However, several experimental design features could have also influenced the present findings, including the timing of PAF measurement, the sequence of stimulation relative to pain induction, and the target site used.

First, the possibility that PAF was modulated but not captured in this study due to the timing of PAF measurement should be considered to explain the present results. Previous research outside the pain field has only observed transient changes in PAF lasting for 1–2 minutes immediately after single 10 Hz rTMS sessions to the pre-frontal cortices (Anderson et al., 2007; Millard et al., 2024; Okamura et al., 2001). Therefore, measuring PAF the day after the final rTMS session in the present study, may explain why effects of rTMS on PAF were not observed. Given the potentially short duration of effects (Anderson et al., 2007; Millard et al., 2024; Okamura et al., 2001), future research should explore both transient PAF alterations immediately after each rTMS session, as well as sustained changes following a greater number of repeated sessions, for example, by matching to the ∼10 rTMS sessions recommended for treatment of chronic pain (Lefaucheur et al., 2020). In addition, the effect of rTMS on PAF during stimulation should be considered, as transient effects might still relate to rTMS-induced pain relief but may not persist beyond the stimulation period. Despite the inherent challenge of measuring oscillations during rTMS due to electrical artifacts (Wagner et al., 2007), doing so could be vital for understanding how rTMS influences pain and the role of PAF in these effects.

Second, the target used for rTMS stimulation could be considered. The DLPFC was chosen based on the above-mentioned systematic review (Millard et al., 2024) that found studies showing changes in PAF after single 10 Hz rTMS sessions to the pre-frontal cortices (Anderson et al., 2007; Okamura et al., 2001) and due to the key role this region plays in descending pain modulation pathways (Glasser et al., 2016; Lorenz et al., 2003; Seminowicz and Moayedi, 2017). However, as PAF from the sensorimotor region has now been linked to pain in several studies including a large cohort with NGF-induced pain (Chowdhury et al., 2025), modulating PAF in the M1 region may be relevant or required to see concurrent changes in PAF and pain.

Third, the focus on concurrent rather than sequential changes in PAF and pain should be considered. Previously reported stability of PAF after NGF injection (Chowdhury et al., 2023; De Martino et al., 2021; Furman et al., 2019) does not necessarily preclude the possibility of NGF-induced pain confounding rTMS-induced effects on PAF. Moreover, assessing concurrent change also hinders the ability to investigate causal relationships due to the absence of a clear temporal sequence (Hayes, 2022). To overcome this, future research could measure the effects of rTMS on PAF before then inducing and measuring experimental pain, to assess whether changes in PAF mediate effects of rTMS on pain. This approach explores potential prophylactic applications of PAF modulation and was further examined in a subsequent study, involving several of the present co-authors, to again investigate rTMS effects on PAF and NGF-induced experimental pain (Chowdhury et al., 2024).

The second study incorporated the above-mentioned modifications to the experimental design, including administering all rTMS sessions before NGF pain induction to separate effects, measuring PAF changes immediately after the final session rather than the following day, and stimulating M1 instead of DLPFC. The findings indicated that, compared to sham, five days of active M1-rTMS increased sensorimotor PAF and was followed by reduced pain ratings. As discussed by Chowdhury et al. (2024), future research should further disentangle how PAF modulation is influenced by target site, stimulation sequence, and timing of PAF measurement.

### 4.2 Closer proximity of PAF to 10 Hz is associated with improved analgesic effects

Although change in PAF was not observed after rTMS in this study, the exploratory finding – that closer absolute proximity of baseline PAF to the stimulation frequency of 10 Hz is associated with greater analgesic effects of rTMS in the active but not sham rTMS group – could provide important information and future direction. This exploratory finding suggests that PAF may influence the analgesic effects of rTMS, perhaps not through its modulation but through individual differences in baseline PAF affecting treatment outcomes. Based on recent literature reviews (Ciampi de Andrade and García-Larrea, 2023; Fernandes et al., 2022), this is, to our knowledge, the first example of the relevance of baseline PAF for the analgesic effects of rTMS. And although this is a novel finding in pain research, it aligns with prior research in depression, where the proximity of baseline PAF to 10 Hz has been associated with enhanced efficacy of rTMS for reducing depressive symptoms (Corlier et al., 2019; Roelofs et al., 2021). For example, a recent study found that patients with major depressive disorder (MDD) and comorbid severe pain were less likely to respond to rTMS treatment, and PAF was positively associated with pain severity (Corlier et al., 2023). Additionally, lower baseline PAF phase-coherence in the somatosensory and default mode networks was observed in patients with comorbid pain who did not respond to treatment (Corlier et al., 2023).

The exploratory nature of the present analysis, coupled with a limited sample size, mandates cautious interpretation. Future research should replicate the potential for baseline PAF to influence the efficacy of rTMS on pain, whilst also considering the effects of rTMS on change in PAF. Nevertheless, the present finding and the findings of previous depression research underscore the potential of PAF as a predictor of rTMS efficacy, suggesting that individual baseline PAF may be crucial factors in treatment response for individuals with both depression and pain. As PAF has been demonstrated to be a clinically accessible measure (Luo et al., 2022; Millard et al., 2022b), a simple screening could be used to identify individuals with PAF close to 10 Hz that could benefit from 10 Hz rTMS treatment. Alternatively, this finding could suggest that rTMS should be applied at each individual’s baseline PAF, to reduce the absolute proximity for all participants. For example, work in depression suggests that synchronising rTMS to individual alpha rhythms could improve clinical outcomes (Pantazatos et al., 2023). Therefore, this exploratory finding opens several new research avenues and is highly relevant given the modest success rates observed in rTMS chronic pain treatment, with only 20–30% of patients reporting significant pain improvement (Attal et al., 2021; Fernandes et al., 2022).

Moreover, recent research suggests that the choice of a fixed 10 Hz stimulation frequency may not be the ideal method for altering PAF. Instead, individualised frequencies based on each participant’s PAF have shown promise (Di Gregorio et al., 2022). Di Gregorio et al. (2022) used rhythmic-TMS with only five pulses in a short burst, with PAF measured between these bursts. Crucially, instead of a fixed 10 Hz frequency, the stimulation frequencies were individualised to be either 1 Hz above or 1 Hz below each participant’s PAF (i.e., PAF *±*1 Hz). This personalised approach is thought to modulate PAF via entrainment, a theory that describes the alignment of neural oscillations to an external rhythmic input (Frohlich and Riddle, 2021; Lakatos et al., 2019; Pikovsky et al., 2001; Thut et al., 2011a). Indeed, Di Gregorio et al. (2022) found that stimulating above individual PAF increased PAF speed, and stimulating below individual PAF decreased PAF speed (Di Gregorio et al., 2022). However, these assessments were conducted during stimulation and are thus subject to stimulation artifacts (Wagner et al., 2007), rather than using a pre-post stimulation design employed in the present study. Note that no studies have assessed the influence of smaller increments in frequencies (e.g., PAF *±*0.2 Hz or PAF *±*0.5 Hz) (Millard et al., 2024), despite evidence for it being easier to entrain internal rhythms that are closer in frequency to external rhythms (Thut et al., 2011a; Vogeti et al., 2022), in line with the physics of synchronization (Pikovsky et al., 2001).

### 4.3 Decreases in fast alpha band power occur in the progression of pain without changes in PAF

Our investigation revealed that there were no discernible effects on alpha power within the broad alpha band (8–12 Hz) or the slow alpha band (8–9.8 Hz) following the administration of NGF-induced pain and rTMS. However, we did observe decreases in fast alpha band power (10–11.8 Hz) for both the active and sham rTMS groups in the progression of pain. This aligns with recent literature that demonstrated decreases in alpha power, one in centro-parietal regions after pain by hot water immersion compared to warm water immersion (Valentini et al., 2022), and another during phasic thermal heat pain and topical capsaicin-heat pain compared to resting state baseline (Furman et al., 2021). Future research is needed to determine whether this change was due to NGF-induced pain or the application of rTMS, regardless of whether the rTMS was active or sham.

While a decrease in fast alpha power might suggest a corresponding shift in PAF due to the CoG calculation method involving the weighted sum of spectral elements across the 8–12 Hz band, our results indicate that substantial changes in fast alpha power can occur without significant impacts on PAF speed. This observation aligns with descriptions in previous analytical research (Chiang et al., 2011, 2008; Furman et al., 2021), underlining that PAF measured by CoG can remain stable even when substantial changes in alpha power occur.

### 4.4 Exploratory Effects of Pain on PAF

In our exploratory analysis, we found no significant correlations between baseline PAF and pain ratings at Day 5 for the entire sample or within individual groups, except for a significant positive correlation between baseline PAF and muscle soreness at Day 5 in the sham group. This positive correlation aligns with findings from some researchers (De Martino et al., 2021; Nir et al., 2010), while it contradicts findings from others (Furman et al., 2020, 2019, 2018; Millard et al., 2023). Additionally, linear mixed-effects models revealed significant interactions between pain intensity and group on PAF. Specifically, while higher pain ratings and muscle soreness were associated with higher PAF in the sham rTMS group, these relationships were less pronounced or possibly negative in the active rTMS group. These findings, although intriguing, should be interpreted with caution due to the small sample size and exploratory nature of this analysis, warranting further investigation in larger cohorts.

### 4.5 Study limitations and constraints

Several limitations warrant consideration. The first limitation concerns the fact that this study only used one pain model (i.e. NGF), whereas incorporating multiple pain models could provide a more comprehensive understanding of the differential effects of interventions on different types of pain, as demonstrated in our other recent work on concurrent changes in PAF and pain (Millard et al., 2023) and recommended for the investigation of the PAF–pain relationship (Mazaheri et al., 2022). Second, individual differences in pain responsiveness could be assessed at baseline to understand their contribution to rTMS efficacy. Third, the potential concurrent and sequential changes in PAF and pain with rTMS could be assessed in chronic and acute clinical pain populations, to complement work conducted in experimental pain.

Lastly, it is crucial to consider PAF not only in the context of pain but also in relation to broader aspects of perception and its potential associations, or lack thereof, with other conditions, responses, and states (Broeke et al., 2013; Zebhauser et al., 2023). For instance, as well as to chronic pain (Zebhauser et al., 2023), PAF has been linked to cognitive functioning (Klimesch, 2012), attention deficit hyperactivity disorder (Arns et al., 2008), and treatment response in depression (Arns et al., 2012; Corlier et al., 2019; Roelofs et al., 2021), whilst its associations with other somatosensory conditions, such as chronic post-burn itch, remain unestablished in the literature (Millard et al., 2022a). Therefore, future studies should continue to explore the presence or absence of a relation to PAF and its modulation in different conditions and settings to better understand its role in health and disease.

### 4.6 Conclusion

Our secondary analysis revealed no significant PAF changes following five days of rTMS, despite previous reports of analgesic effects in this cohort (De Martino et al., 2019; Seminowicz et al., 2018). Nevertheless, exploratory findings indicate that individuals with baseline PAF closer to 10 Hz might be more responsive to rTMS, offering a potential target group for this treatment. Future research should validate these findings in larger samples and expand upon them by assessing the potential of individualised stimulation frequencies to enhance the efficacy of rTMS as a pain management approach.

## 5 Acknowledgements

The authors would like to acknowledge Christian L. Christiansen for their copy editing of this manuscript.

## 6 Abbreviations

ANOVA: Analysis of variances
CI: Confidence interval
CoG: Centre of gravity
DLPFC: Dorsal lateral prefrontal cortex
ECRB: extensor carpi radialis brevis
EEG: Electroencephalography
FIR: finite impulse response
M1: Primary motor cortex
MDD: Major depressive disorder
MEPs: Motor evoked potentials
NGF: Nerve growth factor
NRS: Numerical rating scale
PAF: Peak alpha frequency
PPTs: Pressure pain thresholds
Qs: Questionnaires
rTMS: Repetitive transcranial magnetic stimulation
rMT: Resting motor threshold
SD: Standard deviation
SE: Standard error
SEPs: Sensory evoked potentials
tDCS: Transcranial direct current stimulation.

## 7 Data sharing

Data and code will be shared upon reasonable request to the corresponding author.

